# Reproducibility across single-cell RNA-seq protocols for spatial ordering analysis

**DOI:** 10.1101/764191

**Authors:** Morten Seirup, Li-Fang Chu, Srikumar Sengupta, Ning Leng, Hadley Browder, Kevin Kapadia, Christina M. Shafer, Bret Duffin, Angela L. Elwell, Jennifer M. Bolin, Scott Swanson, Ron Stewart, Christina Kendziorski, James A. Thomson, Rhonda Bacher

**Affiliations:** Molecular and Environmental Toxicology Program, University of Wisconsin Madison, Madison, Wisconsin, United States of America; Morgridge Institute for Research, Madison, Wisconsin, United States of America; Department of Biostatistics and Medical Informatics, University of Wisconsin Madison, Madison, Wisconsin, United States of America; Department of Statistics, University of Florida, Gainesville, Florida, United States of America; Department of Cell & Regenerative Biology, University of Wisconsin School of Medicine and Public Health, Madison, Wisconsin, United States of America; Department of Molecular, Cellular, & Developmental Biology, University of California Santa Barbara, Santa Barbara, California, United States of America; Department of Biostatistics, University of Florida, Gainesville, Florida, United States of America

## Abstract

As newer single-cell protocols generate increasingly more cells at reduced sequencing depths, the value of a higher read depth may be overlooked. Using data from three different single-cell RNA-seq protocols that lend themselves to having either higher read depth (Smart-seq) or many cells (MARS-seq and 10X), we evaluate their ability to recapitulate biological signals in the context of pseudo-spatial reconstruction. Overall, we find gene expression profiles after spatial-reconstruction analysis are highly reproducible between datasets despite being generated by different protocols and using different computational algorithms. While UMI based protocols such as 10X and MARS-seq allow for capturing more cells, Smart-seq’s higher sensitivity and read-depth allows for analysis of lower expressed genes and isoforms. Additionally, we evaluate trade-offs for each protocol by performing subsampling analyses, and find that optimizing the balance between sequencing depth and number of cells within a protocol is important for efficient use of resources. Our analysis emphasizes the importance of selecting a protocol based on the biological questions and features of interest.

## Introduction

Single-cell RNA sequencing (scRNA-seq)^1–5^ is a powerful tool for studying transcriptional differences between individual cells. The innovation of droplet-based techniques^6,7^ and unique molecular identifiers (UMI)^8^ has lowered the cost per cell and pushed the field towards obtaining data from tens of thousands of cells per experiment albeit at a reduced sequencing depth. Recent publications have compared the sensitivity, accuracy, and precision between several scRNA-seq techniques and report that the major trade-off between protocols is sensitivity, which is dependent on read depth^9,10^. With the push for sequencing an ever-increasing number of cells at the expense of read depth per cell, the value of a higher read depth might be overlooked. Here we investigate the reproducibility of biological signals across protocols that naturally lend themselves to generating data on more cells versus higher read depth.

Studies comparing protocols have mainly done so with respect to performance on spike-ins or on technical variability alone^9,10^. Recently, Guo et al.^11^ showed agreement of cell types and signature genes between two platforms used for single-cell RNA-seq – Fluidigm C1 and Drop-seq. However, few studies have examined comparative agreement among protocols for biological inferences beyond clustering and identifying differential gene expression, yet a key question of interest with single-cell data is its ability to reflect temporal or spatial heterogeneity. For cells collected at a given time, the underlying dynamic biological process is reflected in genome-wide differences in gene expression. Computational algorithms that attempt to order cells in pseudo-time or pseudo-space based on variability in gene expression have been developed^4,12,13^, and more than 45 existing algorithms were recently compared^14^. Yet, as far as we know, no comparison of single-cell protocols exists for the question of cell ordering.

Here, our evaluation is in the context of pseudo-spatial reconstruction in which we compared three independently produced scRNA-seq datasets on the mouse liver lobule. We chose to compare protocols on their ability to reflect the spatial patterning of the liver lobule in which the parenchymal cells of the liver, hepatocytes, are organized spatially in a polygonal shape around a central vein (Figure 1A). From the central vein, a gradient of metabolic functions is performed extending to a portal triad at each vertex^15–19^. The gradient of differences in gene expression patterns is referred to as the zonation axis (from periportal (PP) to pericentral (PC))^20^. This coordinated spatial organization provides a particularly interesting application of single-cell techniques. For this study we obtained one dataset using Smart-seq—a full-length protocol, a second dataset using MARS-seq^21^—a UMI and plate based protocol and the third dataset generated using 10X^22^—a UMI and droplet protocol. Although the cell number and read depth differ greatly across datasets, we find high reproducibility of gene expression profiles after spatial-reconstruction analysis. Given the reproducibility and that each protocol naturally lends itself to either producing more cells at a lower sequencing depth or fewer cells at a higher depth, our results demonstrate the importance of carefully evaluating the biological question and features of interest when selecting the appropriate sequencing protocol. In applications focused on lower expressed genes or on genes with high sequence similarity, increased read depth is preferable, whereas a focus on identifying cell types based on more highly expressed genes will benefit from collecting more cells. In an ideal situation a single cell assay would result in thousands of cells that are all sequenced at a high read depth, but technical and financial restrictions make this rarely possible.

**Figure 1.**
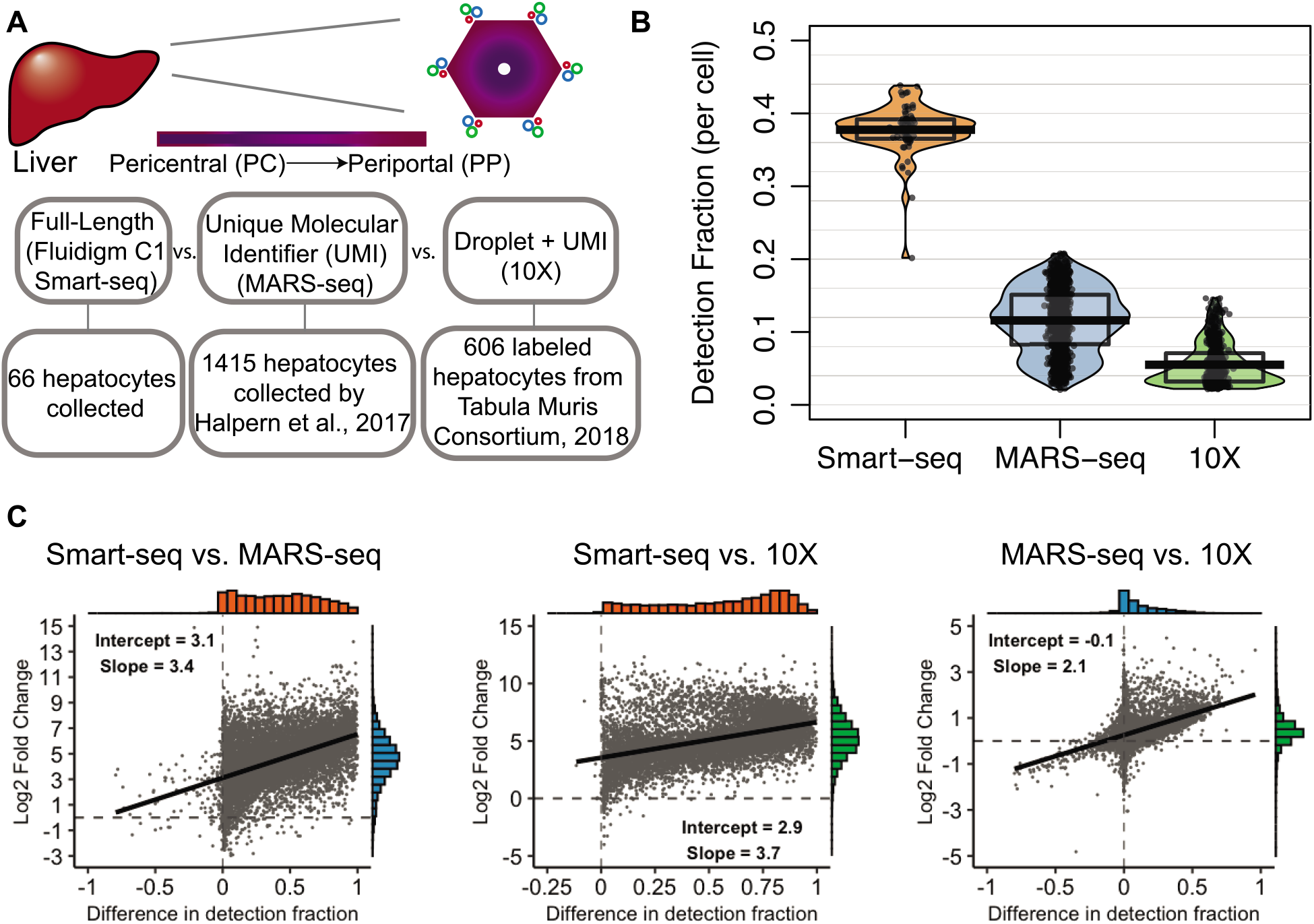
Illustration of the liver anatomy, and general comparison of the datasets. A) Top. Illustration of the liver lobule identifying the portal triad along the outer edges and the central vein in the middle. The color gradient represents metabolic zonation. Bottom. Highlights the main differences between the datasets compared. B) Comparison of gene detection fraction between the datasets. The detection fraction per cell (y-axis) is shown for the two datasets (x-axis). C) Left. The log2 fold-change of genes detected above an average expression level of zero in the Smart-seq dataset compared to the MARS-seq dataset (y-axis), versus the difference in gene-level detection fractions across datasets (x-axis). A linear regression line is overlaid and a histogram of the x- and y-axis are shown opposite of each axis. Middle. Similar plot shown for Smart-seq versus 10X. Right. Similar plot shown for 10X versus MARS-seq.

## Results

### Differences in detection rates

By using the Fluidigm C1 coupled with the Smart-seq protocol, we were able to identify on average around 38% (about 7100 genes) (Figure 1B) of all genes in the genome expressed per cell, whereas the MARS-seq dataset finds on average 12% (about 2200 genes) and the 10X dataset finds on average 6% (about 1100 genes) (Figure 1B). This is in accordance with findings by Ziegenhain et al. 2017 when they examined single-cell transcriptomic methods^9^ and by Phipson et al^23^ when comparing biases in full-length versus UMI protocols. The increased sensitivity of the full-length protocol is further illustrated in Figure 1C, which on a per gene level shows the difference in detection fraction compared to the log fold change in mean expression between the protocols. A difference in detection fraction of zero means that the gene is detected in the same fraction of cells in both datasets. The difference across protocols in log2 fold-change has a linear relationship with the difference in detection fractions, which indicates a fairly constant increase in log2 expression as cells are sequenced with greater sensitivity. At the intercept, a difference in detection equal to zero, the log2 fold change is 3.1 between Smart-seq and MARS-seq, indicating an experiment wide increase in sensitivity in the Smart-seq protocol of approximately 9-fold. Between Smart-seq and 10X, the increase in sensitivity is approximately 12-fold and there is a similar level of sensitivity between MARS-seq and 10X. Not surprisingly, the vast majority of genes are detected in a larger fraction of cells and have a higher expression level in the more deeply sequenced dataset using the Smart-seq protocol. Although, it is worth pointing out that around 6% of genes have higher detection using the MARS-seq protocol (negative values on x-axis) and a few of these genes also have higher expression levels (negative values on y-axis) than in the Smart-seq protocol. This subset of genes better detected in the MARS-seq dataset have higher GC content and are slightly longer (S1 Figure), which is consistent with previous reports of protocol comparisons^23,24^.

### Reconstructing spatial profiles of liver zonation profiles

Next, to represent the spatial patterns across the liver lobule, the cells in the three datasets were computationally ordered according to their expression profiles. The MARS-seq dataset was spatially ordered by Halpern et al. 2017 by first performing smFISH for six marker genes at various locations across the zonation axis, then single-cell RNA-seq data obtained by MARS-seq assigned cells to one of nine zonation locations based on each cell’s expression profile of the six marker genes^21^. We ordered the cells in the 10X dataset using the Monocle2 algorithm, which builds a trajectory through cells based on the expression similarity among the most highly variable genes^12^. For the Smart-seq protocol we used the computational algorithm Wave-Crest to spatially order cells based on fifteen marker genes known in the literature to be differentially expressed along the zonation axis (Figure 2A)^5^. The ordering procedure uses the nearest insertion algorithm implemented in the Wave-Crest package, which searches among the space of all possible orderings via a 2-opt algorithm by considering insertion events and choosing orders which minimize the mean square error of a polynomial regression on the marker genes expression. Of the 15 genes used, we selected eight periportal expressed genes and seven pericentral expressed genes^20^. All orderings assume the zonation profile and spatial organization can be represented in a single dimension. A similar reconstructed order was obtained for the Smart-seq dataset when applying Monocle2 (S2 Figure).

**Figure 2.**
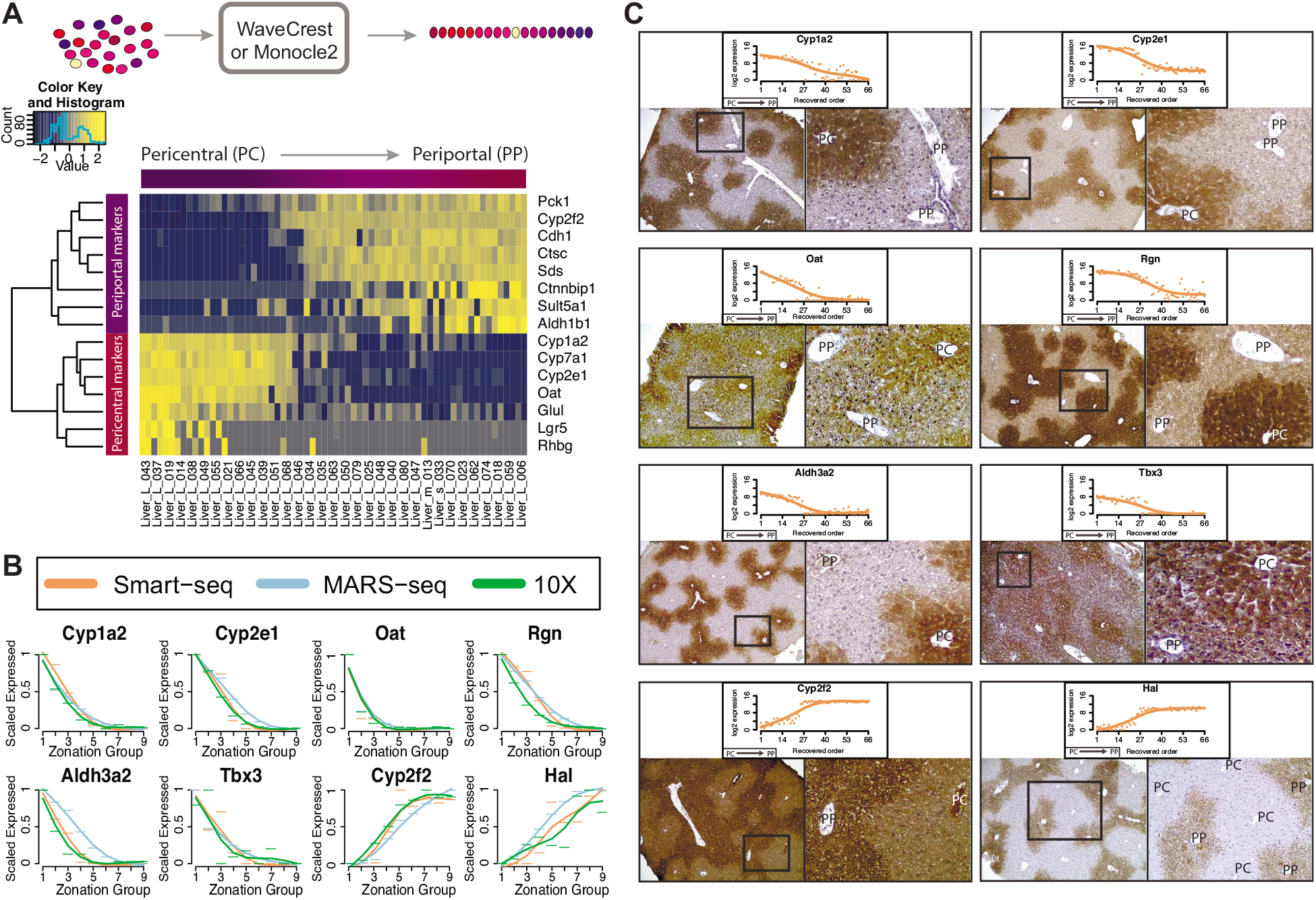
Pseudo-space reordering of hepatocytes, and prediction and validation of dynamically expressed genes. A) Top. Illustration of the pseudo-spatial reordering process. Bottom. Heatmap showing the pseudo-spatial reordering (x-axis) and the expression levels of the marker genes (y-axis) for the Smart-seq dataset. Pericentral cells are found on the left-hand side and Periportal cells are found on the right-hand side. B) Scaled expression profile (y-axis) of 8 dynamic genes based on the predicted pseudo-space reordering (x-axis) of the Smart-seq dataset (orange), the MARS-seq dataset (blue), and the 10X dataset (green). C) Immunohistochemistry staining of the genes highlighted in B). Above the staining is the log2 expression counts (y-axis) across the predicted pseudo-spatial order (x-axis) of the Smart-seq dataset. The left picture shows the staining and the right picture is an enlarged section (black square). PP = Periportal, PC = Pericentral.

Using the recreated order of the hepatocytes, we explored the dynamics of gene expression across the periportal to pericentral axis. Figure 2B shows a subset of genes that are predicted to be highly regulated across the axis, four of which were not in our list of marker genes. Since the MARS-seq dataset placed cells into nine discrete zones along the axis, we divided cells from the Smart-seq and 10X datasets into nine equally sized groups in order to compare the reconstructed orderings. The zonation profiles in Figure 2B have high agreement, with a median correlation of 0.95 between the three datasets. Before proceeding, we also performed an additional experiment to validate that our cell ordering and expression profiles reflect those of the liver lobule *in vivo*. Remarkably, immunohistochemistry studies showed that selected marker gene protein expression profiles also agreed with our spatial reconstructed scRNA-seq datasets: six markers display a PC-high/PP-low profile and two markers display a PC-low/PP-high profile in mouse liver lobule *in vivo* (Figure 2C). This confirmation in protein gradient patterns corresponding to our reconstructed mRNA profiles provides us with confidence for further analysis on the biological inference in comparing the three protocols in this context.

### Comparing marker gene expression across liver zonation profiles

An exciting prospect of single cell analysis is the identification of genes that have non-monotonic or dynamic expression across pseudo-time or space. Several genes in the bile acid synthesis pathway were shown by Halpern et al., 2017 to be non-monotonically expressed in a pattern where the highest expression levels along the lobule correspond to the functional placement of the genes in the bile acid synthesis pathway (Cyp7a1, Hsd3b7, Cyp8b1, Cyp27a1 and Baat)^21^. We find that the expression profiles for these genes are corroborated across the three datasets (Figure 3).

**Figure 3.**
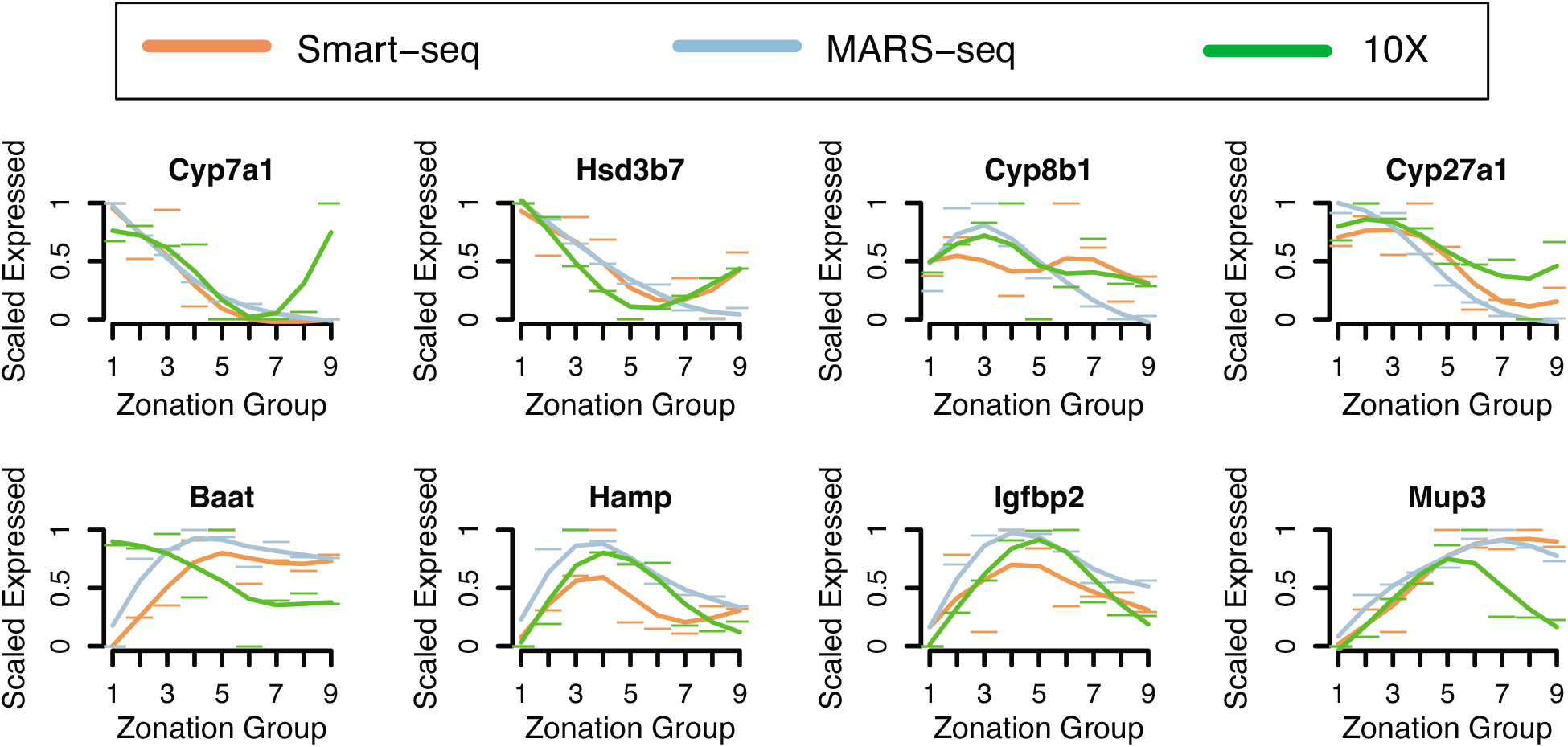
Comparison of zonation profiles across three datasets. Scaled expression profile (y-axis) of 8 genes non-monotonically expressed from Halpern et al. along the predicted pseudo-space reordering (x-axis) of the Smart-seq dataset (orange), the MARS-seq dataset (blue), and the 10X dataset (green).

However, in the Smart-seq dataset, Cyp8b1 is found to have largely flat expression levels along most of the lobule and lower expression toward the periportal zone and Baat appears to have an opposite trend in the 10X dataset. Other genes shown to be non-monotonically expressed such as Hamp, Igfbp2 and Mup3 in Halpern et al., 2017 display similar non-monotonic expression profiles in the Smart-seq and 10X datasets (Figure 3). The ability to identify gene expression profiles that are either high at the PP end, high at the PC end, or high in the middle of the liver lobule confirms that the sampling depth is sufficient to spatially reconstruct the liver lobule. We also investigated the expression pattern of Glul in more detail as it is known to be expressed highly in a one hepatocyte-wide band around the central vein^25^. Accordingly, the predicted expression pattern found using all datasets demonstrated sufficient sampling of this region (S3 Figure).

We further compared the zonation profiles between datasets by identifying genes having significant differential expression along the reconstructed spatial order across the periportal to pericentral axis. For genes displaying differential zonation in all datasets (having adjusted p-value < .1), the Smart-seq versus MARS-seq dataset had the highest median correlation (0.86), while the Smart-seq versus 10X had the lowest median correlation (0.69). In Figure 4A we looked at significantly zonated genes within the metabolic pathways in KEGG and found the median correlation between all datasets ranged from 0.75 to 0.89. When all genes were considered the median correlation ranged from 0 – 0.04.

**Figure 4.**
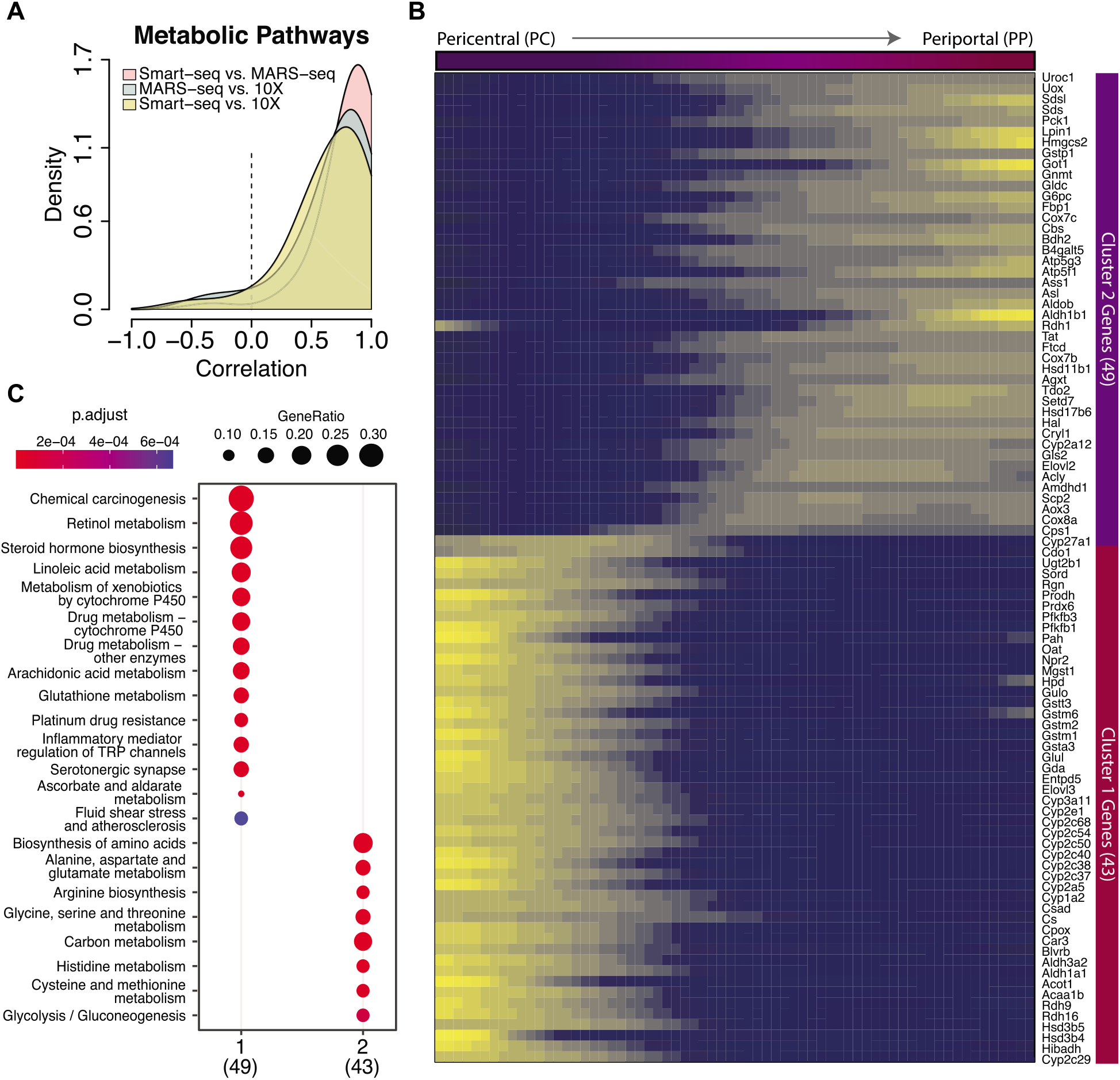
Correlation and Gene Ontology analysis of genes between datasets. A) Correlation analysis of significantly zonated genes annotated to the metabolic pathways in KEGG between the datasets. The pairwise correlation is shown for each dataset comparison. B) Heatmap of the expression level of genes that are significantly differentially zonated in all datasets and enriched in the metabolic KEGG pathway. C) Breakdown of KEGG enrichment analysis of the two k-mean clusters based on the genes shown in B. Dot size represents the fraction of enriched genes in each ontology, and the color represents the adjusted p-value for the enrichment.

Traditionally the liver lobule is divided into three zones, a periportal zone 1, a pericentral zone 3, and transitioning zone 2^26,27^. The transitional nature of the liver axis is reflected in the heatmap of metabolic genes that were significantly zonated in all datasets (Figure 4B). Using k-means clustering, we found the Smart-seq data tended to cluster into two distinct gene groups representing either the periportal or pericentral zone. Examination of the two clusters by enrichment analysis of KEGG metabolic pathways (Figure 4C) revealed that the predicted location along our reconstructed axis of metabolic processes with known periportal or pericentral bias such as amino acid metabolism (periportal), lipogenesis (pericentral), and CYP450 metabolism (pericentral) corresponds to their known *in vivo* locations^27^. Despite using different reordering algorithms and protocols, the three datasets show high agreement of expression along the recovered pericentral to periportal axis among genes that are significantly zonated in all datasets, and reliably mirror the *in vivo* patterning of the liver lobule (additional KEGG categories are shown in S4 Figure).

### Differences in gene profiles among lowly expressed genes and gene isoforms

When we look at genes with moderate and low expression levels, we find that the datasets differ to a greater degree. We identified twenty-one genes that were classified as significantly zonated along the periportal to pericentral axis in the Smart-seq dataset that were not detected at all in the MARS-seq dataset and thirty-five such genes not detected in the 10X dataset. Compared to the Smart-seq dataset, ten genes were exclusively detected in the MARS-seq dataset and no genes were exclusive to the 10X dataset. Figure 5A shows the six most highly expressed genes that we were able to exclusively identify in the Smart-seq dataset having significant zonation (adjusted p-value < .1).

**Figure 5.**
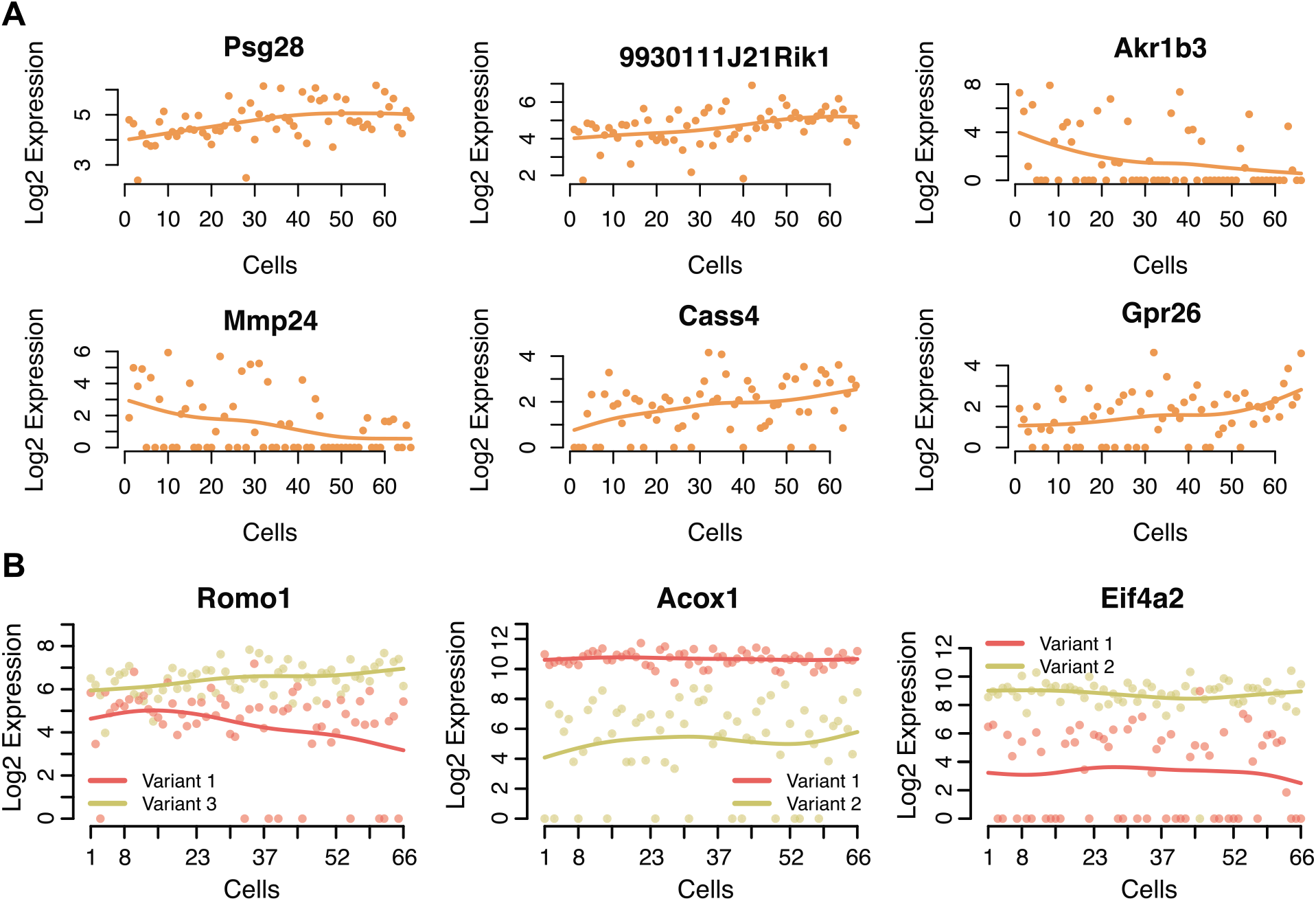
Genes and isoforms found in the full-length dataset and not in the UMI datasets. A) Six genes found to be zonally expressed in the Smart-seq dataset that were not detected in either the MARS-seq or 10X datasets. The log2 of expression values are represented on the y-axis and the pseudo-space ordered cells are found on the x-axis. B) Examples of genes with two transcript variants expressed differently across reordered cells from the Smart-seq dataset.

Further, an exciting field of study that benefits from an enhanced resolution of scRNA-seq is isoform analysis^28–30^. Many genes in the genome have two or more isoforms that are distinctly expressed and can change properties such as structure, function, and localization of the resulting protein^31^. Due to the increased sensitivity of full-length cDNA libraries generated by Smart-seq protocol, we were able to examine genes with known isoforms and identify cases where the transcript variants for each isoform has distinct expression from each other across the periportal to pericentral axis, which is not possible with less sensitive protocols. In Figure 5B the transcript variants of Romo1 are seen to display opposite trends in expression across the zonation axis, where the Romo1 variant 3 is increasing in expression from the pericentral end towards the periportal end and the Romo1 variant 1 is decreasing in expression along the same axis. We also highlight genes Acox1 and Eif4a2 whose variants both show constant expression across the zonation axis but at different levels. Both of these genes are known to have isoform-specific expression in the liver lobule^32,33^. (For Ensembl and ENTEREZ IDs for transcript variants see S6 Table).

Due to UMI based protocols capturing only one end of the transcript compared to full-length cDNA procedures, there is an inability to resolve not just isoforms but also many genes that are closely related. For instance, there were 242 concatenated genes in the MARS-seq set that correspond to 539 unique genes. An example of this is seen in S5 Figure where we highlight a concatenate of Ugt1a enzymes. Eight genes are concatenated (annotated together) and when combined, the average expression level is shown to be high at the pericentral end of the lobule and low at the periportal end. Again, it is clear that not all the members of this concatenated group follow this trend as Ugt1a6a can be seen to have consistent expression levels across the pericentral to periportal axis.

### Evaluating the trade-off within each protocol *in silico*

To further study the trade-offs between higher depth versus more cells, we performed a subsampling experiment. For each dataset, we held either the number of cells or the sequencing depth constant while varying the other. For the Smart-seq and 10X datasets, we evaluated the effect on the cell ordering as well as the gene-specific zonation profiles. For the MARS-seq dataset, the assignment of each cell to a zonation group depended on external data and was independent of the other cells profiled, and thus we only evaluated the effect on zonation profiles. We estimate the MSE (mean squared error) as the difference in zonation profiles in the subsampled dataset versus the original dataset. In Figure 6A, the MARS-seq dataset displayed an approximately linear tradeoff in zonation profile error for fewer cells at the original read depth. However, at reduced read depths using the original 1,415 cells, the error increased exponentially (Fig.6B). Within a dataset, we can compare the MSE between the two trade-off scenarios and we find that for the MARS-seq dataset resequencing at the same depth results in error that is equivalent to the reduction observed in MSE by going from 600 to 1400 total cells. For the 10X dataset, we also find an approximately linear tradeoff in zonation profile error for fewer cells at the original read depth (Fig. 6C). However, at reduced read depths using the original 606 cells, we observe a gradual increase in error as total depth decreases (Fig.6D). Similarly, by comparing the MSE trade-off, it appears that resequencing at the same depth results in error that is equivalent to reducing the total cells from 600 to around 400. Thus, in scenarios with very low sequencing depth (average of 3-12k total UMIs per cell), sequencing deeper may be more beneficial than adding more cells. For the Smart-seq dataset, we found the spatial ordering to be quite robust to reduced sequencing depth, even as low as 50% fewer reads only marginal increased the average MSE as shown in Figure 6F. The average sequencing depth for the Smart-seq cells was 3.5 million counts per cell, well beyond the suggested sequencing saturation for single-cell data that occurs close to one million total reads^34^. We do see more dramatic increases in error related to zonation profiles when profiling fewer cells (Figure 6E). For Smart-seq data, sequencing to even half of the current depth and increasing the number of cells would be beneficial.

**Figure 6.**
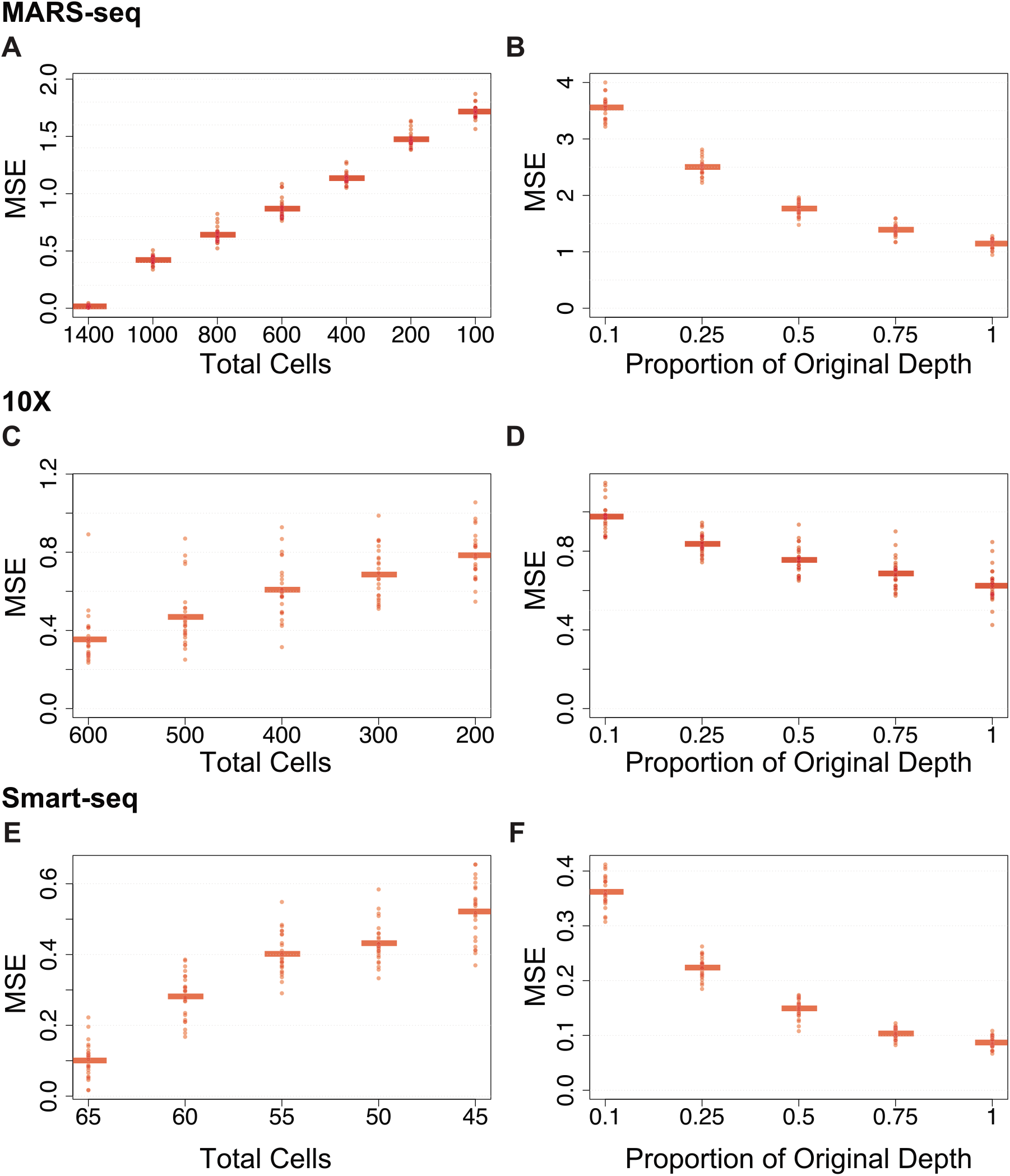
Subsampling total numbers of cells and sequencing depth. A) For 25 subsamplings at various total numbers of cells in the MARS-seq dataset, the mean squared error (MSE) of the zonation profile over 500 randomly selected genes is shown. B) Similar to A, but for 25 subsamplings at various total read depths. C-D) Similar to A-B, but for the Smart-seq dataset. E-F) Similar to A-B, but for the 10X dataset.

## Discussion

In summary, we compared three scRNA-seq datasets of mouse hepatocytes where two, MARS-seq and 10X, are wide but shallow and the other, Smart-seq, is narrow but deeply sequenced. We find that the three different protocols present highly reproducible liver zonation profiles in single cells, and for the vast majority of genes that are highly expressed we observe highly comparable results. Our results were not dependent on any one computational method or pre-processing pipeline. We do however find that when we look at medium to low expressed genes, the increased sensitivity of the C1/Smart-seq protocol is able to identify several genes exclusive to this dataset. This increased sensitivity also allowed us to identify several genes with isoforms that behaved differently across the periportal to pericentral axis. Though in general, there are still limitations of short reads in regard to isoform analysis and if more accuracy is needed, the newly developed technique ScISOr-seq^35^ might be better suited. We do however believe that this full-length data allows for more reliable preliminary isoform analysis compared to either UMI method. However, the main weakness of using fewer cells is that it is less likely that rare cell types will be sampled. In cases where such rare cells are of high interest, protocols that produce a large number of cells are preferable. In an ideal case, one would sample many cells and sequence all of them deeply; unfortunately, this is not always possible in practice and the decision of whether to sample many cells shallowly or fewer cells deeply comes down to whether rare cell types are of interest or if higher resolution of the individual cells is preferred.

Given the distinct advantages of the protocols, we emphasize that the biological question should be the driving factor when deciding on protocol. Within a chosen protocol, achieving balance between the sequencing depth and the number of cells is still an important consideration for optimal use of resources. Based on our simulations of datasets at opposite ends of the sequencing depth versus number of cells trade-off, there is eventually a detriment to sacrificing reads for additional cells or sequencing beyond the attainable sensitivity level on too few cells. We expect that the extent of the cells versus depth trade-off will vary for other cell types or tissues and it will largely depend on the heterogeneity of the biological system under study.

## Methods

### Animals and handling

All animals were kept under standard husbandry conditions. A wildtype 8-week-old male C57BL/6 (Jackson laboratories) was used in this experiment. Using isoflurane, the mouse was anesthetized before euthanizing by cervical dislocation. Animal experiments and procedures were approved by the University of Wisconsin Medical School’s Animal Care and Use Committee and conducted in accordance with the Animal Welfare Act and Health Research Extension Act.

### Cell isolation

The euthanized mouse was pinned to a Styrofoam plate using 20 ga needles to aid in dissection. The abdominal cavity was opened, and the portal vein exposed. A piece of 4-0 suture thread (Ethicon vicryl coated) was threaded under the portal vein and used to secure a 26 ga catheter inserted into the portal vein (Butler Schein animal health 26 G IV Catheter, Fisher Scientific). Hepatocytes were isolated using a 2-step perfusion protocol. First, Liver Perfusion Medium (Gibco) warmed to 37°C was pumped through the catheter for 10 minutes using a peristaltic pump at 7 ml/min flowrate. Then, Liver Digest Medium (Gibco) warmed to 37°C was pumped through the liver at the same settings for 10 minutes. After perfusion, the liver was excised and transferred to a 10 cm dish containing 20 ml liver digest medium. The liver was dissected, allowing the cells to spill into the media. The cells were then filtered through a 40 μm cell strainer into a 50 ml tube and 30 ml media (Williams E media + 2 μg/ml human insulin + 1x glutamax + 10% FBS) were added and placed on ice. The hepatocytes were purified by centrifugation at 50 x G, 4 times for 3 minutes each, each time discarding the supernatant and adding media.

### Single cell RNA sequencing-Full-length dataset

Single-cell RNA sequencing was performed as previously described^4,5^ with the following modifications. In this study, we used small (5-10 μm), medium (10-17 μm), and large (17-25 μm) plate sizes. ERCC RNA Spike-In (ThermoFisher Cat. No. 4456740) was diluted in the lysis mix following the manufacturer’s user guide and previous studies^36^. Single end reads of 51 bp were sequenced on an Illumina HiSeq 2500 system.

Sequencer outputs were processed using Illumina’s CASAVA-1.8.2. The demultiplexed reads were trimmed and filtered to eliminate adapter sequence and low-quality basecalls. The reads were mapped to an mm10 mRNA transcript reference (extended with ERCC transcripts) using bowtie-0.12.9^37^; expression estimates were generated using RSEM v.1.2.3^38^. Using the Fluidigm C1 system to capture and synthesize cDNA from single cells in the liver, we generated transcriptomes for 149 cells. To exclude low quality transcriptomes, we removed cells in which the fraction of ERCC spike-in made up 20% or more of the total assigned reads. This left 66 high quality cells that were used in the downstream analysis. Finally, the data was normalized using SCnorm (R package v 1.5.7)^39^.

### Pseudo-spatial reordering-Full-length dataset

For the full-length data, the cells were computationally ordered using the Wave-Crest method as described in Chu et al. 2016^5^. For the reordering step, gene expression values were rescaled to mean 0 and variance 1 to ensure the values across different genes are comparable. The Wave-Crest algorithm implements an extended nearest insertion algorithm that iteratively adds cells to the order and selects the insertion location as the location producing the smallest mean squared error in a linear regression of the proposed order versus gene expression. A 2-opt algorithm is then used to find an optimal cell order by considering adjacent cell exchanges. The cell ordering step uses the expression profiles of pre-selected known marker genes of liver zonation. Thus, the resulting linear profile of ordered cells represents the periportal to pericentral axis. The known marker genes used to construct the periportal to pericentral axis in Wave-Crest include the following pericentral markers: cytochrome P450 7a1 (Cyp7a1), cytochrome P450 2e1 (Cyp2e1), ornithine aminotransferase (Oat), cytochrome P450 1a2 (Cyp1a2), rh family, B glycoprotein (Rhbg), leucine-rich repeat-containing G-protein coupled receptor 5 (Lgr5), glutamate-ammonia ligase (Glul); and the following periportal markers: phosphoenolpyruvate carboxykinase 1 (Pck1), catenin beta interacting protein 1 (Ctnnbip1), aldehyde dehydrogenase 1 family member B1 (Aldh1b1), sulfotransferase family 5A, member 1 (Sult5a1), cytochrome P450 2f2 (Cyp2f2), cathepsin C (Ctsc), serine dehydratase (Sds), and E-cadherin (Cdh1). All markers were selected based on their expression ratio as reported by Braeuning et al. 2006^20^.

A detection step was done to identify additional genes that follow the one-dimensional periportal to pericentral axis by fitting a linear regression to the relationship between each gene’s expression and the Wave-Crest cell order. To determine if a gene is significantly dynamic (differentially zonated) along the recovered axis, we tested whether the regression slope is different from zero. We reported the Benjamini-Hochberg adjusted p-values to control the false discovery rate. For genes having an adjusted p-value < .01, the direction of the expression profile was assigned based on the sign of the regression slope (periportal: positive slope, pericentral: negative slope). We also calculated the linear fitting mean squared error (MSE) for each gene. Genes with a smoother trend over the recovered cell order are expected to have a smaller MSE. We report the full list of genes, sorted by their MSE, in S7 Table; scatter plots for genes having adjusted p-value < .01 are shown in S8 File.

### Pseudo-spatial reordering-10X dataset

The 10X dataset was downloaded from the Tabula Muris compendium public resource via Figshare^22^. The 10X data was originally processed using the CellRanger version 2.0.1. Within the liver cells, the authors originally identified 975 hepatocytes. For our analysis, we performed a second quality control step to identify cells with low RNA content, possible doublets, or dead/damaged cells, where we filtered cells based on the total number of genes expressed per cell. Using the Seurat R package v3.1.5, hepatocytes were further filtered to those having between 200 and 3000 genes detected per cell (only one cell had more than 5000 genes detected per cell). Next, we clustered the cells using Seurat, where a k-nearest neighbors (KNN) graph used was constructed based on the first 20 principle components to create a shared nearest neighbors graph based on the Jaccard index between each cell and its 20 nearest neighbors, as implemented in the FindNeighbors function. Clusters were then identified by partitioning this graph using the Louvain community detection algorithm with a resolution of .5, as implemented in the FindClusters function. The cells clustered into three distinct larger groups and we retained only the largest grouping of cells that clustered together, resulting in 606 total cells. The data was then normalized using scran v1.12.1. Next, we used Monocle v2.12.0 to order the cells, basing the ordering on the top 200 highly variable genes estimated using the mean variance relationship via the FindVariableFeatures function in Seurat. To determine if a gene is significantly dynamic (differentially zonated) along the recovered axis, the Monocle2 function differentialGeneTest was used to fit a spline on gene expression versus the estimated pseudo-time.

### Comparative Analysis

Smoothed densities (bean plots) with overlaid raw data, the mean, and a box representing the interquartile range of the cellular detection fractions were created using the pirateplot function in the yarrr R package (v0.1.5). The cellular detection fraction was calculated per cell as the proportion of genes having expression greater than zero. The fold-change for each gene between the two datasets (A versus B) was calculated as the log2 fold-change of the dataset A over dataset B, where each gene mean was calculated as the average expression among non-zero counts across all cells in the datasets. The heatmap in Figure 2 of marker gene expression on the normalized Smart-seq data was generated by setting values above the 95th percentile or below the 5nd percentile to the 95th percentile or 5nd percentile value, respectively.

Due to the datasets having different dynamic ranges, we used scaled expression plots to compare expression profiles, where the ordered cells in the full-length dataset and 10X were each divided into nine equally sized groups to correspond to the nine layers in the UMI dataset. For the full-length and 10X dataset, for a given gene, the median (full-length) or mean (10X) expression in each group was calculated, then the nine values were scaled between zero and one. Smoothed fits were overlaid using the smooth.spline function in R with the degrees of freedom parameter df=4. Expression correlations along the zonation axis between datasets were calculated using Pearson correlation. Enrichment of genes in KEGG pathways or GO was done using the R package clusterProfiler (v. 3.10.1)^40^. For the enrichment analysis, since different statistical methods were used to assess zonation profiles, genes were considered significantly zonated if they had an adjusted p-value < .1 in all datasets. The heatmap in Figure 3 is a smoothed heatmap, where a smoothing spline was first fit to the log expression (pseudo-count of one added) of each gene using the smooth.spline function in R with the smoothing parameter df=4 which provided profiles that were not over- or underfit in either dataset. Then the smoothed expression was scaled and outliers above the 98^th^ percentile or below the 2^nd^ percentile were set to the 98^th^ percentile or 2^nd^ percentile value, respectively. Additional KEGG categories from this analysis can be interactively viewed on Github https://github.com/rhondabacher/scSpatialReconstructCompare-Paper.

### Subsampling Analysis

In all subsamplings described below, each scenario was repeated a total of 25 times and the zonation group means were scaled to be between zero and one.

For the MARS-seq dataset, zonation group means were recalculated on a subsampled set of cells using the posterior probability matrix and original UMI counts from Halpern et al. 2017. In each sampling, the mean squared error (MSE) was calculated based on a random sample of 500 genes as 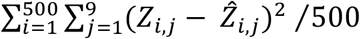, where *Z_i,j_* represents the mean expression of gene *i* in zonation group *j* in the original dataset and 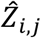 is the corresponding value for the subsampled dataset. For subsampling at lower read depths, we fixed the number of cells at the original total of 1415 cells and simulated each cell’s gene counts individually using a multinomial distribution. For each cell, the subsampled total counts were set to X% of the original total read counts for that cell (for X = (10,20,30,40,50,60,70,80,90,100)) and each gene’s cell-specific probability was calculated as its original count divided by the original total counts for that cell. The MSE was calculated for each subsampled set as described above.

For the Smart-seq dataset, we reran Wave-Crest when subsampling the total number of cells using the original parameter settings and marker genes. Then, as before, the ordered cells were assigned zonation groups by dividing cells into nine equally sized groups. The zonation profile error was estimated using MSE and calculated as described above with the exception that since Wave-Crest orders can be flipped, we calculated the MSE on the returned order and its reverse, and kept the minimum MSE of the two. To evaluate the zonation profile error with lower read depths, we used a similar approach as described above for the MARS-seq dataset, fixing the number of cells to be the same as the original total of 66 and, since the order correlation was shown to be consistently high, we used the original Wave-Crest order for every scenario when evaluating zonation profile error. For the 10X dataset, the subsampling was performed similarly as for the Smart-seq dataset, however Monocle2’s ordering was more variable as it was not based on marker genes and thus we did not fix the order when evaluating the zonation profile error. Trade-offs in MSE are directly comparable within a dataset but due to intrinsic differences in the original processing and in subsampling, the MSE should not be compared across the datasets.

### Immunohistochemistry

An 8-week-old male C57BL/6 mouse was anesthetized using isoflurane before euthanizing by cervical dislocation. The liver was excised, sliced as thinly as possible with a razor blade, and fixed in formaldehyde overnight. The liver slices were paraffin embedded and sectioned. Sections were stained following the protocol published by Abcam (http://www.abcam.com/ps/pdf/protocols/ihc_p.pdf). In short, the slices are deparaffinized by dipping into sequential solutions of 100% xylene, 50-50% xylene-ethanol, 100% ethanol, 95% ethanol, 70% ethanol, 50% ethanol, and tap water. The antigens were then retrieved by placing the slides in Tris-EDTA buffer (10 mM Tris Base, 1 mM EDTA Solution, 0.05% Tween 20, pH 9.0) and incubating them in a decloaking chamber (Biocare Medical Decloaking Chamber #DC2008US) with the following settings: delayed start 30 sec.; preheat 80°C, 2 min.; heat 101°C, 3 min. 30 sec.; and fan on. The slides were washed 2 x 5 min in TBS + 0.025% Triton X-100 before they were blocked for two hours at room temperature in 10% normal serum in 1% BSA. The appropriate primary antibody was then diluted in the same 10% normal serum in 1% BSA, added to the slides, and incubated at 4°C overnight in an incubation chamber. The next day the slides were washed 2 x 5 min in TBS + 0.025% Triton X-100 followed by 15 min incubation in 0.3% H2O2 at room temperature. Next, the appropriate secondary antibody was diluted into 10% normal serum in 1% BSA before it was added to the slides and incubated for 1 hour at room temperature. The slides were then washed 3 x 5 min in TBS before DAB (#ab103723) staining mixed according to manufacturer instruction was applied and incubated under a microscope to stop the reaction after sufficient staining. The slides were rinsed in tap water for 5 min before being counterstained with Mayer’s hematoxylin (#MHS1-100ML) for 30 sec. The stain was developed in running tap water for 5 min. The slides were then dehydrated by sequentially dipping in 50% ethanol, 70% ethanol, 95% ethanol, 100% ethanol, 50-50% xylene-ethanol, and 100% xylene before Poly-Mount (#08381-120) was added and a coverslip placed on top. The following primary antibodies were added: Aldh3a4 1:250 (AB184171), Cyp2e1 1:50 (AB28146), Cyp1a2 1:50 (R31007), Rgn 1:100 (NBP1-80849), Oat 1:50 (AB137679), Cyp2f2 1:100 (SC-67283), Hal 1:50 (AV45694), and Tbx3 1:50 (SC-31657). The following secondary antibodies were used: goat-anti-rabbit HRP conjugated (ab97051) and donkey-anti-goat HRP conjugated (ab97110) at a concentration of 1:500.

## Supporting information

All Supplemental Figures

## Acknowledgements

Not applicable.

## Supporting information captions

S1 Figure – Examining GC content and gene length in genes with a higher detection fraction in either dataset. Top) The GC content (left) and gene length (right) are shown for genes having a higher detection fraction in either the Smart-seq dataset (gray) or the MARS-seq dataset (blue). A dotted line is shown for genes having a larger mean in either dataset. The two lines closely correspond since the genes having a high detection fraction typically have a higher mean. Bottom) Similar to the top for comparing the Smart-seq and 10X datasets.

S2 Figure – Correlation between WaveCrest and Monocle methods for ordering cells in the Smart-seq dataset.

S3 Figure – Expression of Glul. Scaled expression plots of Glul showing high correlation among all three datasets.

S4 Figure – Correlation analysis of more KEGG pathways. A) Top left: Correlation analysis for genes in the KEGG pathway “Complement and coagulation cascade”. The pairwise correlation is shown for each dataset comparison. Following are plots for the eight highest correlated genes between the any two datasets in that pathway. On the right is a smoothed heatmap of the Smart-seq expression data for the gene expression of all significantly zonated genes enriched in that KEGG pathway. B) Similar to (A) but for the “Drug metabolism – cytochrome P450” pathway. C) Similar to (A) but for the “Biosynthesis of amino acids” pathway.

S5 Figure – Additional genes in Smart-seq dataset but not in the MARS-seq dataset. Eight Ugt1a genes that were concatenated in the MARS-seq dataset (blue on all graphs), but can be resolved in the Smart-seq dataset (orange line).

S6 Table – Ensembl and RefSeq ID’s for genes with transcript variants.

S7 Table – Summary of genes with dynamic expression across the zonation axis identified using Wave-Crest.

S8 File – Scatter plots of dynamic genes listed in S6 Table.

S9 Dataset – Normalized Smart-Seq single-cell data with cells in the Wave-Crest order.

