## Supplementary figures and images for "Reproducibility across single-cell RNA-seq protocols for spatial ordering analysis"

### All Supplemental Figures

#### S1 Figure

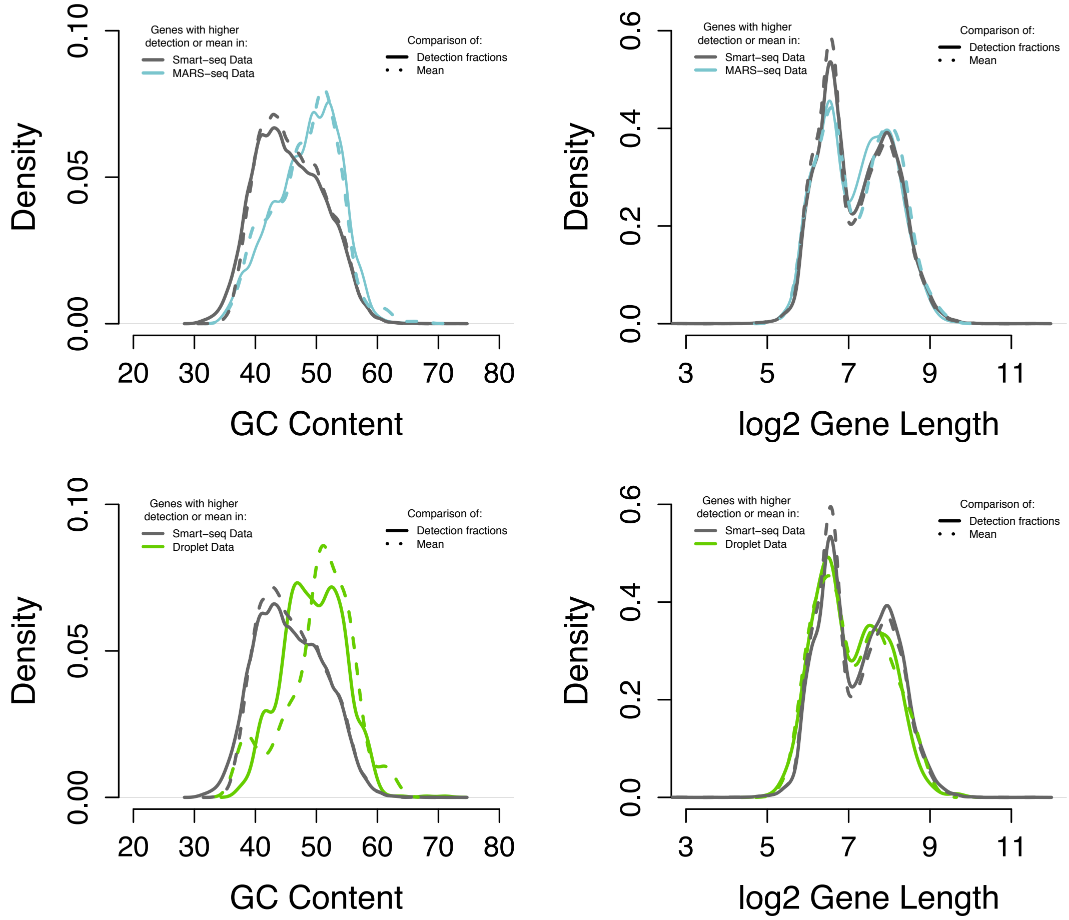


#### S2 Figure

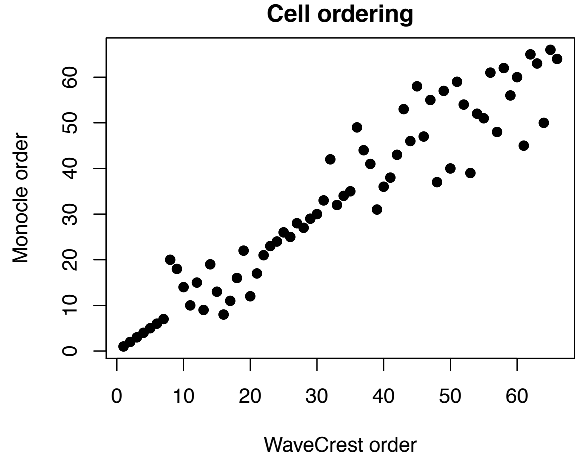

**S3 Figure**

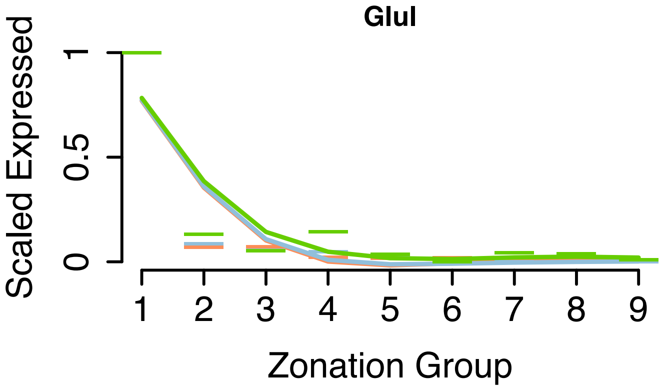

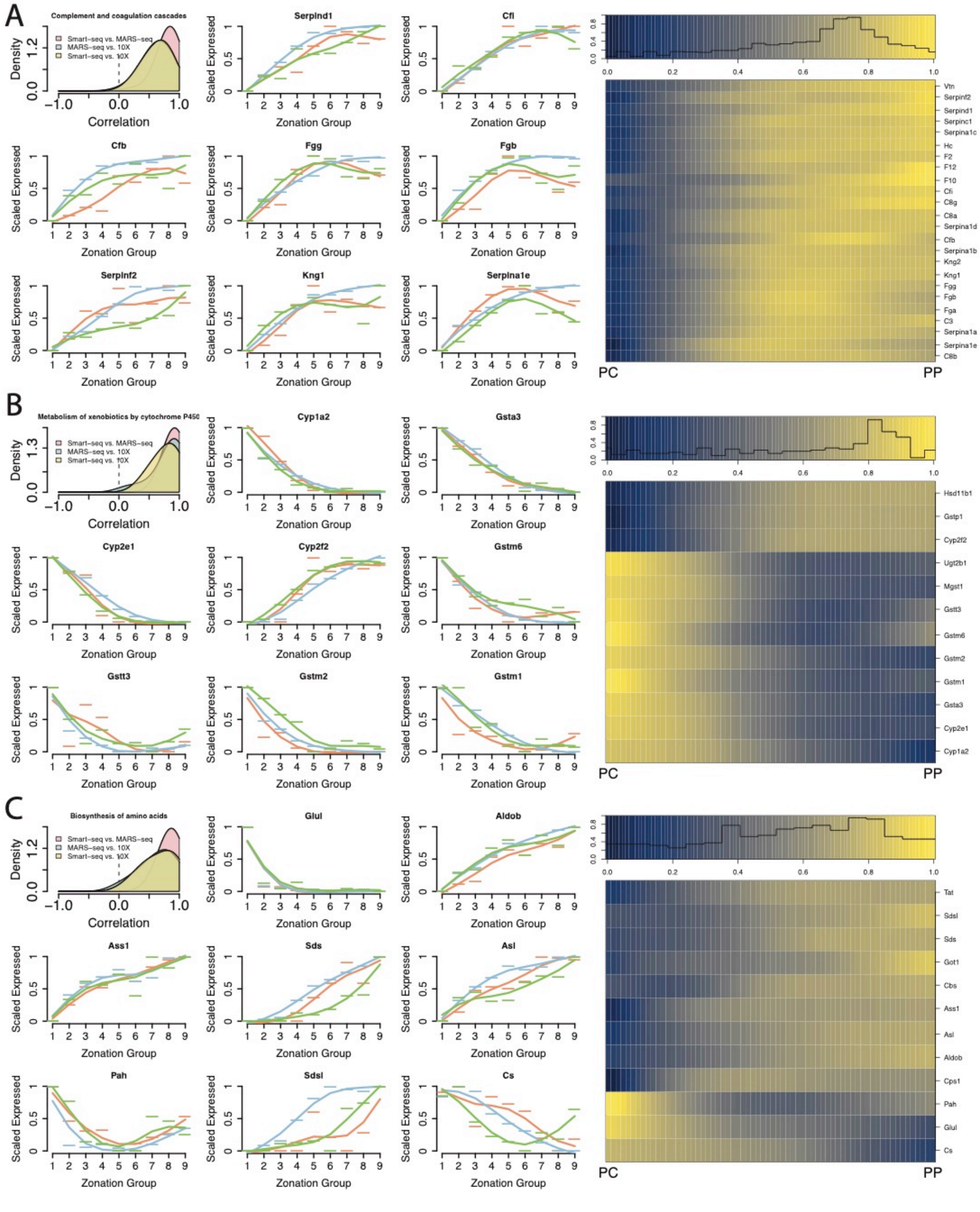

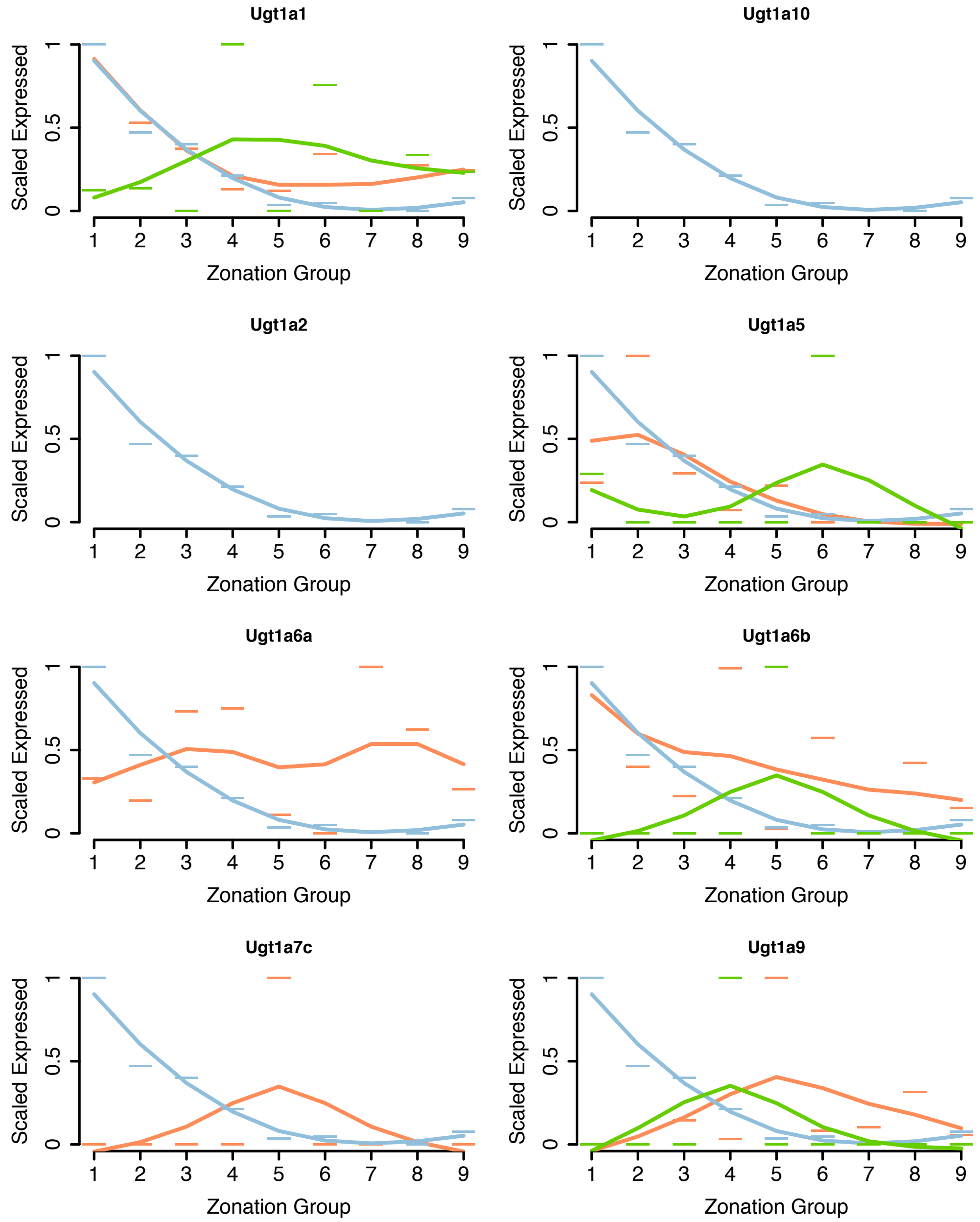
